# Automated Detection of Ripple Oscillations in Long-Term Scalp EEG from Patients with Infantile Spasms

**DOI:** 10.1101/2020.06.03.132183

**Authors:** Colin M. McCrimmon, Aliza Riba, Cristal Garner, Amy L. Maser, Daniel W. Shrey, Beth A. Lopour

## Abstract

**Objective:** Scalp high frequency oscillations (HFOs) are a promising biomarker of epileptogenicity in infantile spasms (IS) and many other epilepsy syndromes, but prior studies have relied on visual analysis of short segments of data due to the prevalence of artifacts in EEG. Therefore, we set out to develop a fully automated method of HFO detection that can be applied to large datasets, and we sought to robustly characterize the rate and spatial distribution of HFOs in IS.

**Methods:** We prospectively collected long-term scalp EEG data from 13 subjects with IS and 18 healthy controls. For patients with IS, recording began prior to diagnosis and continued through initiation of treatment with adenocorticotropic hormone (ACTH). The median analyzable EEG duration was 18.2 hours for controls and 83.9 hours for IS subjects (∼1300 hours total). Ripples (80-250 Hz) were detected in all EEG data using an automated algorithm.

**Results:** HFO rates were substantially higher in patients with IS compared to controls. In IS patients, HFO rates were higher during sleep compared to wakefulness (median 5.5/min and 2.9/min, respectively; *p* =0.002); controls did not exhibit a difference in HFO rate between sleep and wakefulness (median 0.98/min and 0.82/min, respectively). Spatially, the difference between IS patients and controls was most salient in the central/posterior parasaggital region, where very few HFOs were detected in controls. In IS subjects, ACTH therapy significantly decreased the rate of HFOs.

**Discussion:** Here we show for the first time that a fully automated algorithm can be used to detect HFOs in long-term scalp EEG, and the results are accurate enough to clearly discriminate healthy subjects from those with IS. We also provide a detailed characterization of the spatial distribution and rates of HFOs associated with infantile spasms, which may have relevance for diagnosis and assessment of treatment response.

## Introduction

Infantile spasms (IS) are associated with developmental delay and an electroencephalographic (EEG) pattern known as hypsarrhythmia, and this triad of symptoms is referred to as West syndrome [1]. IS carries a poor prognosis, as spasms may evolve into persistent neurological deficits, intractable seizures, and Lennox-Gastaut syndrome, and they carry an increased mortality risk [2, 3]. Not only are these outcomes devastating to patients’ families, but they also put a tremendous burden on the already-strained U.S. healthcare system [3]. Because early treatment initiation may positively impact outcomes [3–5], prompt diagnosis is crucial. However, IS and ictal events may not be captured without prolonged video-EEG monitoring, and the classically associated interictal EEG pattern known as hypsarrhythmia, characterized by high amplitude asynchronous slow waves with multifocal spikes and polyspikes [6], may not be present [6–8] or may have highly variable features [9, 10], leading to poor inter-rater reliability [11, 12]. Thus discovery of other epileptic spasms-associated EEG features is imperative to facilitate prompt and accurate diagnosis.

One such feature may be interictal cortical high-frequency oscillations (HFOs). HFOs are electrographic events above 80 Hz, corresponding to oscillations in the ripple band (80-200 Hz) and fast ripple band (200-500 Hz), as well as to the upper end of the gamma band (∼60-200 Hz) [13]. In healthy individuals, the information encoded in these frequencies are thought to play a role in diverse cortical processes, such as sensory processing, memory consolidation, attention, and movement planning/control [14–17]. In patients with epilepsy, HFOs are thought to be temporally associated with ictal events [18, 19] and are also spatially associated with seizure onset zones [20, 21]. In fact, even interictal HFOs may spatially correlate with seizure onset zones [22–24] and may be useful in predicting long-term outcomes of epilepsy surgery [25].

Many studies support that EEG is a reliable, noninvasive modality to study scalp HFOs in the ripple [26–29] and fast ripple frequency bands [30–32]. Studies using simultaneous scalp and intracranial EEG have confirmed that scalp-visible oscillations have neural origin [29–31]. Scalp HFOs have been associated with various pediatric epilepsies, including benign epilepsy with centrotemporal spikes [33], epileptic encephalopathy with continuous spike-and-wave during sleep [34, 35], idiopathic partial epilepsy of childhood [36], tuberous sclerosis complex [32], and myoclonic seizures [37]. Visual detection is standard in such studies, due to the pervasive nature of scalp EEG artifacts. However, semi-automated detection methods have been proposed, in which candidate HFOs are identified via an automated algorithm and all detections are visually reviewed to reject false positives [29, 32, 33, 38]. von Ellenrieder et al. (2012) created a fully automated detection method, but they recommended expert review of detected events [28]. Our group also published a fully automated method for detection of scalp HFOs, but it was applied to a small dataset (∼10 minutes of data per subject) [39].

HFOs in the gamma and ripple bands have also been reported during epileptic spasms using ictal scalp EEG [18–20, 40, 41]. In some cases, the HFOs were found to be focal, despite the appearance of a generalized spasm onset [19, 40]. Fewer studies have examined HFOs in IS using interictal data. One study [38] analyzed both ictal and interictal HFOs in patients with epileptic spasms without focal features in hypsarrhythmia, and they compared the results to scalp EEG from normal control subjects. They found higher rates of interictal HFOs in the gamma and ripple frequency bands in patients with West Syndrome, and the occurrence of HFOs decreased following treatment with ACTH. A second study found that beta and gamma scalp oscillations were more frequently associated with IS compared to other pediatric epilepsies [22]. However, these studies only utilized a few minutes of interictal data for each subject, given the time-intensive nature of visually inspecting HFOs, and thus the generalizability of their findings remains unclear.

Therefore, we sought to further characterize interictal HFOs in patients with epileptic spasms using long-term recordings and fully automated methods. We measured scalp EEG in normal controls and patients with IS, performed automatic detection of HFOs, and compared the following: 1) the rates of HFOs during wakefulness and sleep, 2) the spatial distribution of HFOs across the cortex, and 3) the effect of treatment initiation on HFO rate.

## Methods

### Subject Recruitment

This prospective longitudinal study was approved by the Children’s Hospital of Orange County (CHOC) Institutional Review Board. Children aged 0-3 years who were admitted to the hospital for long-term video EEG monitoring with concern for new onset epileptic spasms were approached for informed parental consent. Subjects with a previous diagnosis of epileptic spasms and those who had been previously treated with adrenocorticotropic hormone (ACTH) or vigabatrin (VGB) were excluded from the study. It should be emphasized that long-term video EEG monitoring was performed solely based on clinical suspicion for epileptic spasms and was in no way influenced by our study. The developmental status of each subject was evaluated by a board-certified neuropsychologist (Amy Maser, PhD) using the Vineland Adaptive Behavior Scales, Third Edition [42].

### Video-EEG Monitoring

Subjects underwent continuous video EEG monitoring while being recorded with 19 scalp electrodes placed according to the international 10-20 system [43] and two mastoid reference electrodes. Data from these electrodes were simultaneously acquired at 5 kHz for the research study and at 200 Hz for clinical monitoring, using a Neurofax EEG-1200 acquisition system with JE-120A amplifier fitted with a QI-124A dual data stream recording unit (Nihon Kohden, Tokyo, Japan). Electrooculographic (EOG), electromyographic (EMG), and electrocardiographic (ECG) data were also acquired at 5 kHz and 200 Hz using the same system. The initial recording duration for each subject was overnight (approximately 18-24 hours), at which time a board-certified pediatric epileptologist reviewed the video-EEG data for clinical evidence of epileptic spasms. Control subjects were defined as those who had (1) no known neurological diseases, (2) no abnormal neuroimaging, (3) normal video EEG recordings with the events of concern captured, and (4) normal scores on the Vineland-3. Subjects with evidence of ES who were started on ACTH (first-line therapy) were assigned to the spasms group and continued to undergo continuous video-EEG monitoring for an additional 48-96 hours using the aforementioned study and clinical data acquisition parameters. Spasms group subjects were designated as “responder” if they exhibited cessation of clinical spasms and their EEG no longer met criteria for hypsarrhythmia (defined as high amplitude, (>200 microvolt) continuous arrhythmic delta activity with multifocal independent epileptiform discharges and lack of normal background activity) during a follow-up overnight video EEG recording performed approximately 2 weeks after their initial ES diagnosis was made. “Non-responders” had persistent spasms and/or hypsarrhythmia at follow-up.

### Analysis

All study data were processed using the same custom MATLAB (The MathWorks, Natick, MA) software. This software: 1) removed amplifier saturation (clipping) artifacts from the dataset, 2) re-referenced the remaining signals to a longitudinal bipolar montage [44], 3) detected HFOs as described in [45]. Briefly, HFO detection involved applying an infinite impulse response (IIR) band-pass filter (80-250 Hz, 10th order) and rectifier to the data, and then identifying any high-amplitude oscillations, i.e. *≥*3 consecutive oscillations with amplitude greater than the upper-limit of background high-gamma activity (defined as the gamma cumulative distribution function from the surrounding 5s of data with *α*=0.0001). While finite impulse response (FIR) filters are typically preferred, IIR filters drastically reduce computation time and have been used successfully for HFO analysis [46]. We chose the IIR filter settings by visually confirming the similarity to data filtered with the equivalent FIR filter. Any HFOs with corresponding electrode “pop” artifacts [47] (defined as DC shifts >50 *µ*V) or muscle artifacts (defined as line length >1900 with units 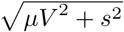 within a 0.3-second sliding window from the longitudinal-bipolar-referenced signals were discarded. Note that the voltage and line length cutoffs were empirically determined from a very small subset of the data by visually inspecting the candidate HFOs detected as above and manually modifying the cutoffs to reject apparent “pop” and muscle artifacts while preserving true HFOs. The subsequent optimal cutoffs were then applied to the remaining dataset.

#### HFO Rates

To compare HFOs in spasms subjects versus control subjects, the total number of HFOs per minute summed across all electrodes was calculated for each subject. Note that in this analysis, to avoid overcounting HFOs with long durations or single HFOs captured by multiple channels (e.g. via volume conduction), any HFO that occurred <1 second after a previous HFO (even from a different channel) was discarded. Manual EEG sleep staging was performed for all control subject studies by a registered polysomnographic technologist (Cristal Garner, REEGT, RPSGT) in accordance with the American Academy of Sleep Medicine (AASM) guidelines. Manual sleep staging was not possible for IS subjects given the size of the datasets and the unreliability of manual sleep staging in this patient population due to epileptic spasms and hypsarrhythmia altering sleep stage structure and progression [48–51]. However, by generating a classifier from EOG, EMG, and ECG data in the control subjects, we were able to mark periods of sleep and wakefulness in IS subjects solely from these non-EEG signals that are likely just as reliable in IS subjects as in control subjects. Briefly, this classifier utilized linear discriminant analysis [52] for dimensionality reduction of the EOG, EMG, and ECG data, followed by a Bayesian classifier using a Gaussian likelihood model. A Wilcoxon rank-sum test [53] was used to compare the HFO rates between control subjects and IS subjects during wakefulness and during sleep. A Wilcoxon signed-rank test for paired data [54] was used to compare the HFO rates during wakefulness and sleep for control and IS subjects.

#### Spatial Distribution of HFOs

To compare the spatial distribution of HFOs throughout the scalp in control and IS subjects, we calculated the number of HFOs per minute at each electrode for every subject during wakefulness and sleep. Note that unlike above, HFOs that occurred <1 second after a previous HFO from a different channel were not discarded in this analysis. This ensured that the spatial distribution of each event was accurately represented; if an event occurred simultaneously in two channels, we counted the event for both channels, rather than arbitrarily removing it from one of them. Lastly, to compare the distributions across subjects, the HFO rate from each electrode was normalized by dividing by the sum of HFO rates across all electrodes for that subject.

#### HFO Rates Before and After ACTH Initiation

A Wilcoxon signed-rank test was used to compare HFO rates prior to and >24 hours after initiating ACTH therapy in IS subjects while awake and asleep. Note that each pair consisted of one patient’s HFO rate during either wakefulness or sleep (summed across all electrodes while avoiding overcounting identical to the *HFO Rates* section above), and the two samples of EEG were clipped as follows: 1) from the start of video-EEG monitoring to the first dose of ACTH, and 2) from 24 hours after the first dose of ACTH to the completion of video-EEG monitoring.

## Results

### Subject Enrollment/Video-EEG Monitoring

Thirty-one subjects were enrolled in the study and contributed ∼1300 total hours of analyzable EEG data. Eighteen subjects were assigned to the control group (4 males, median age 6.7 months, lower quartile 3.0 months, upper quartile 8.2 months). Median useable recording duration was 18.2 hours (lower quartile 16.3 hours, upper quartile 20.9 hours). For all control subjects, the events of interest were captured on video EEG and found to not resemble epileptic spasms. All of the control subjects’ EEG studies were within normal limits for age with no slowing, epileptiform discharges, nor seizures noted. Only one control subject (subject 4) had neuroimaging (MRI) performed, and it was within normal limits. All control subjects had Vineland-3 developmental scores within normal limits (greater than 85). The classifier that used EOG, EMG, and ECG data to predict wake and sleep states for IS subjects was generated from the control subjects’ data and had a mean leave-one-out cross-validation accuracy of 86% when compared with the manual sleep staging results for control subjects. Note that EKG data from control subjects 12 and 17 had significant artifacts requiring additional processing steps before they were used to generate the wake versus sleep classifier.

Thirteen of the thirty-one enrolled subjects demonstrated epileptic spasms (spasms group) during video-EEG monitoring (4 males, median age 7.5 months, lower quartile 6.9 months, upper quartile 14.3 months). Median recording duration was 83.9 hours (lower quartile 41.8 hours, upper quartile 110.4 hours). Clinical data for these subjects are shown in Table 1. This group included individuals with genetic syndromes (trisomy 21, CDKL5, 2q24 and 2p24 deletion, 5q deletion) as well as those with a history of occipital lobe epilepsy, intracranial hemorrhage w/ gliosis, and sinus venous thrombosis. IS subject 1 was on phenobarbital and levetiracetam at the start of the study. IS subject 4 was on phenobarbital at the start of the study. IS subject 7’s EKG and EMG data had significant artifacts requiring additional processing steps to distinguish between wakefulness and sleep. IS subject 9 was initially diagnosed with focal seizures and was started on Phenobarbital, however, clinical follow-up with a subsequent video EEG captured epileptic spasms as well, and the patient was started on ACTH therapy 1 month after the initial video-EEG monitoring. Due to technical difficulties, only ∼2 hrs of data was recorded from IS subject 10 at the study-specified acquisition rate (5 kHz) before the subject started ACTH therapy. Note that the clinical recording was unaffected. Also note that subject 10 was prescribed levetiracetam at the study onset. IS subject 12 only participated in the initial ∼24 hrs of video-EEG monitoring and was started on ACTH therapy immediately after. IS subject 13 was prematurely withdrawn from the study by her parents after only 3 hours and 40 minutes of EEG monitoring; her data are included only in Table 1 below. Still, in total, ∼1300 hours of EEG data were analyzed, and >900,000 HFOs were detected.

**Table 1.**
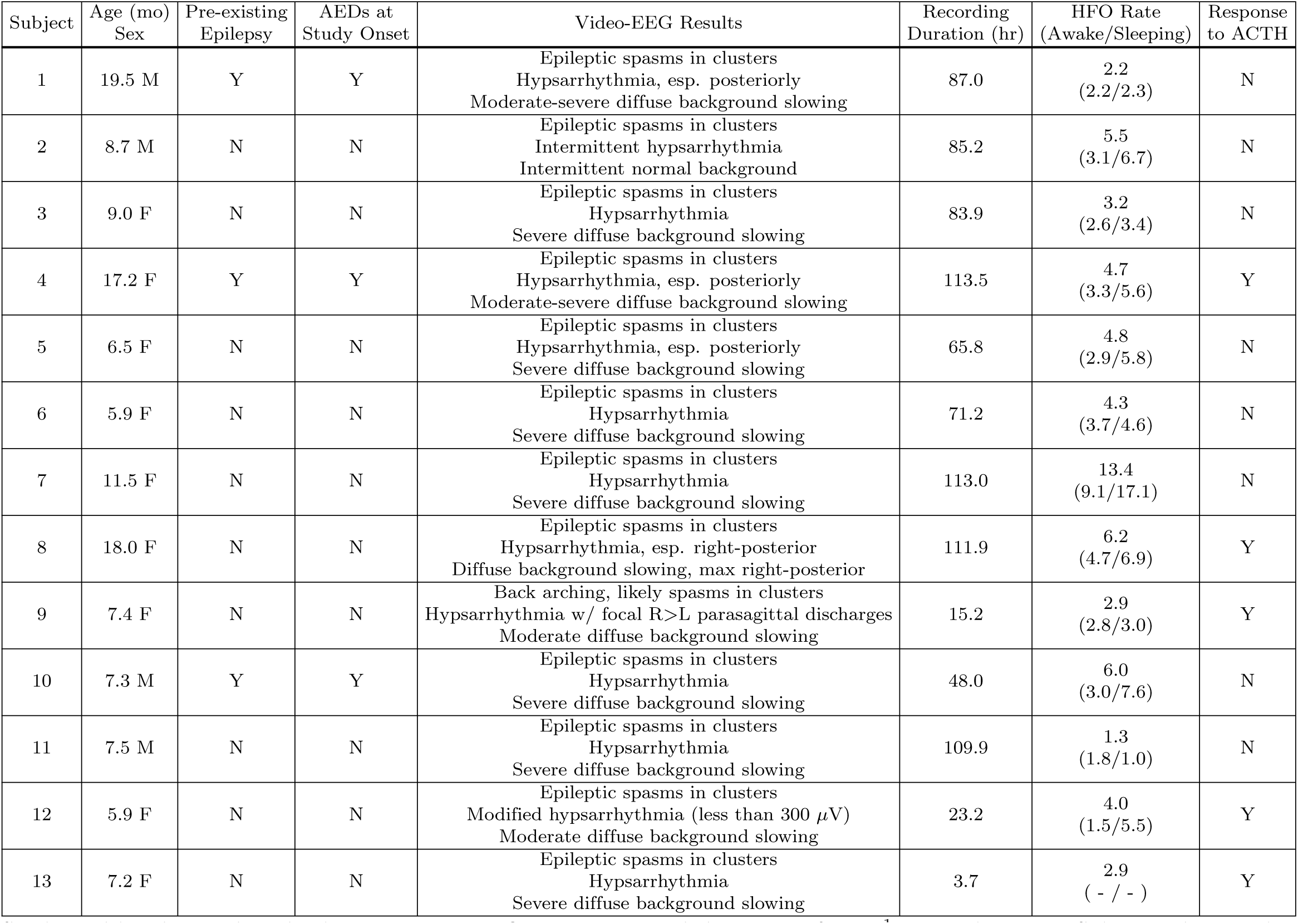
IS subjects’ baseline and study characteristics. HFO rates are provided in units of min^−1^. Note that given Subject 13’s very short recording duration, HFO rates were not calculated separately for wakefulness and sleep. AEDs = antiepileptic drugs, ACTH = adrenocorticotropic hormone.

### Analysis

#### HFO Rates

As depicted in Figure 1, the median HFO rate for the control subjects was 0.82/min while awake and 0.98/min while asleep, and the median HFO rate for IS subjects was 2.9/min while awake and 5.5/min while asleep. HFO rates during both wakefulness and sleep were significantly increased in IS subjects compared to control subjects (*p <*0.0001), per a Wilcoxon rank-sum test. For IS subjects, there was a significant increase in the HFO rate while asleep compared to while awake (*p* =0.0024), per a Wilcoxon signed-rank test for paired data. For control subjects, there was no significant difference between the HFO rates while asleep and while awake (*p* =0.20).

**Fig 1.**
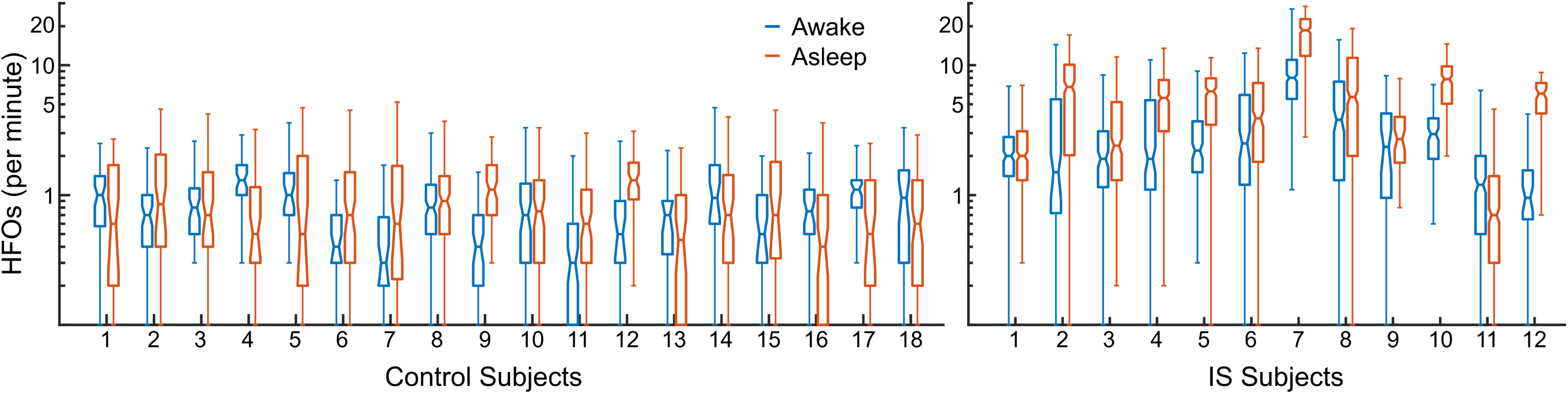
HFO Rates Across Control and IS Subjects. HFOs were counted within multiple 10-minute intervals, and the distribution of HFO rates (number of HFOs per minute across all channels) from these 10-minute intervals is depicted here for control subjects and IS subjects while awake and asleep. Note that a logarithmic y-axis scale is used for the box-and-whisker plots (interquartile range within box and all values within whiskers) given the discrepancy in HFO rates between the control and IS subjects.

#### Spatial Distribution of HFOs

As depicted in Figures 2 and 3, HFOs in the control subjects typically occurred outside of the posterior parasagittal region corresponding to bipolar signals from C3-P3, P3-O1, Fz-Cz, Cz-Pz, P4-O2, C4-P4 (the center colum of channels outlined in these Figures). The HFO rates had an otherwise broad distribution both while awake and during sleep, although the trend is more striking during sleep. In contrast, most of the HFOs in IS subjects during wakefulness and sleep were concentrated in a few channels, and these channels tended to include rather than spare the posterior parasagittal region. These comparisons can be seen clearly in Figure 4, which depicts the median spatial distribution of HFOs across control subjects and IS subjects while awake and asleep.

**Fig 2.**
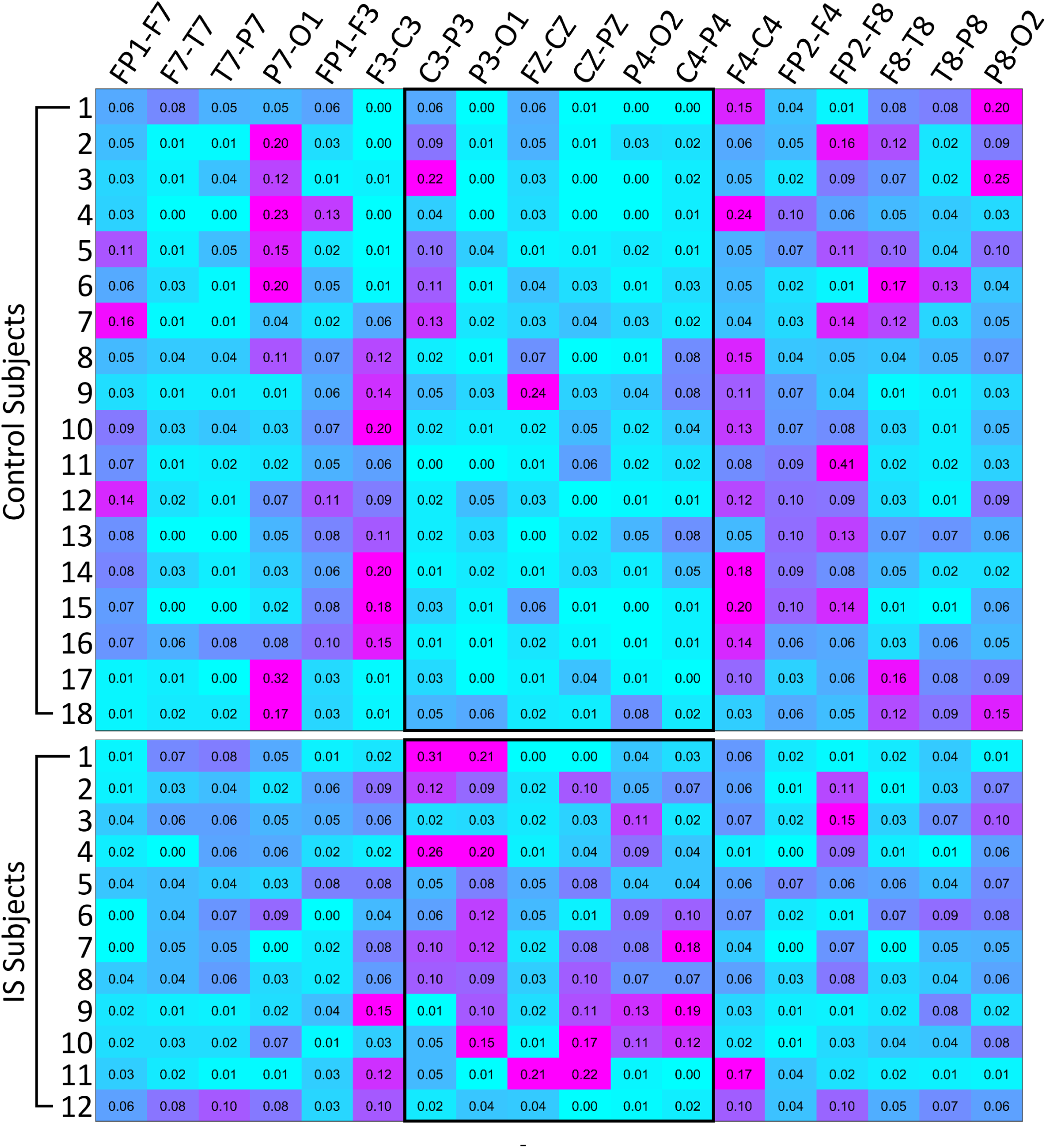
Normalized HFO Rates for Each Channel During Wakefulness. Each row represents a control or IS subject and each column corresponds to a channel from a longitudinal bipolar montage. HFO rates are normalized across each subject such that the sum of each row is 1.0, and higher rates appear pink while lower rates appear cyan. The channels contained in the black outlines correspond to posterior parasagittal areas.

**Fig 3.**
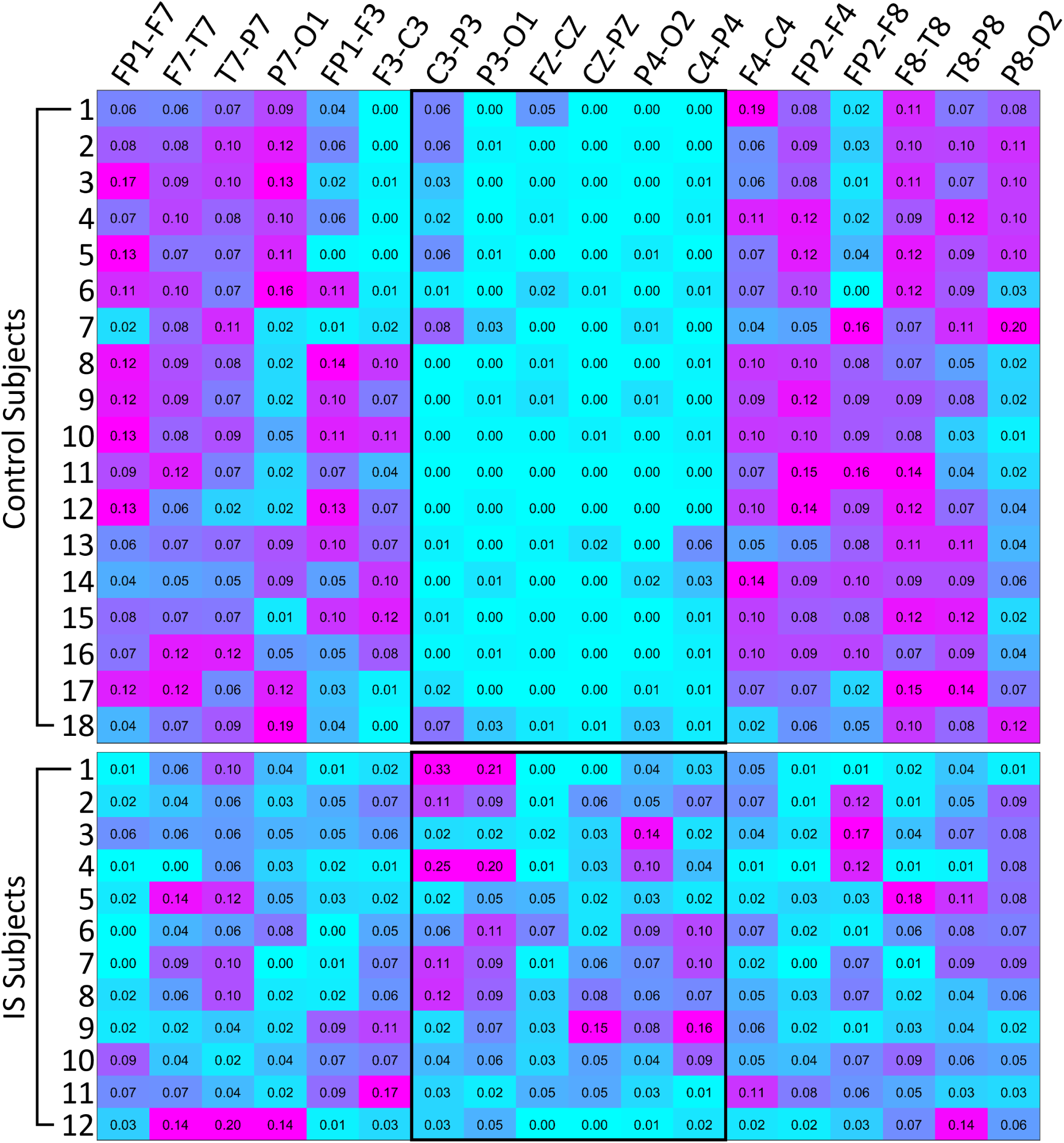
Normalized HFO Rates for Each Channel During Sleep. Each row represents a control or IS subject and each column corresponds to a channel from a longitudinal bipolar montage. HFO rates are normalized across each subject such that the sum of each row is 1.0, and higher rates appear pink while lower rates appear cyan. The channels contained in the black outlines correspond to posterior parasagittal areas.

**Fig 4.**
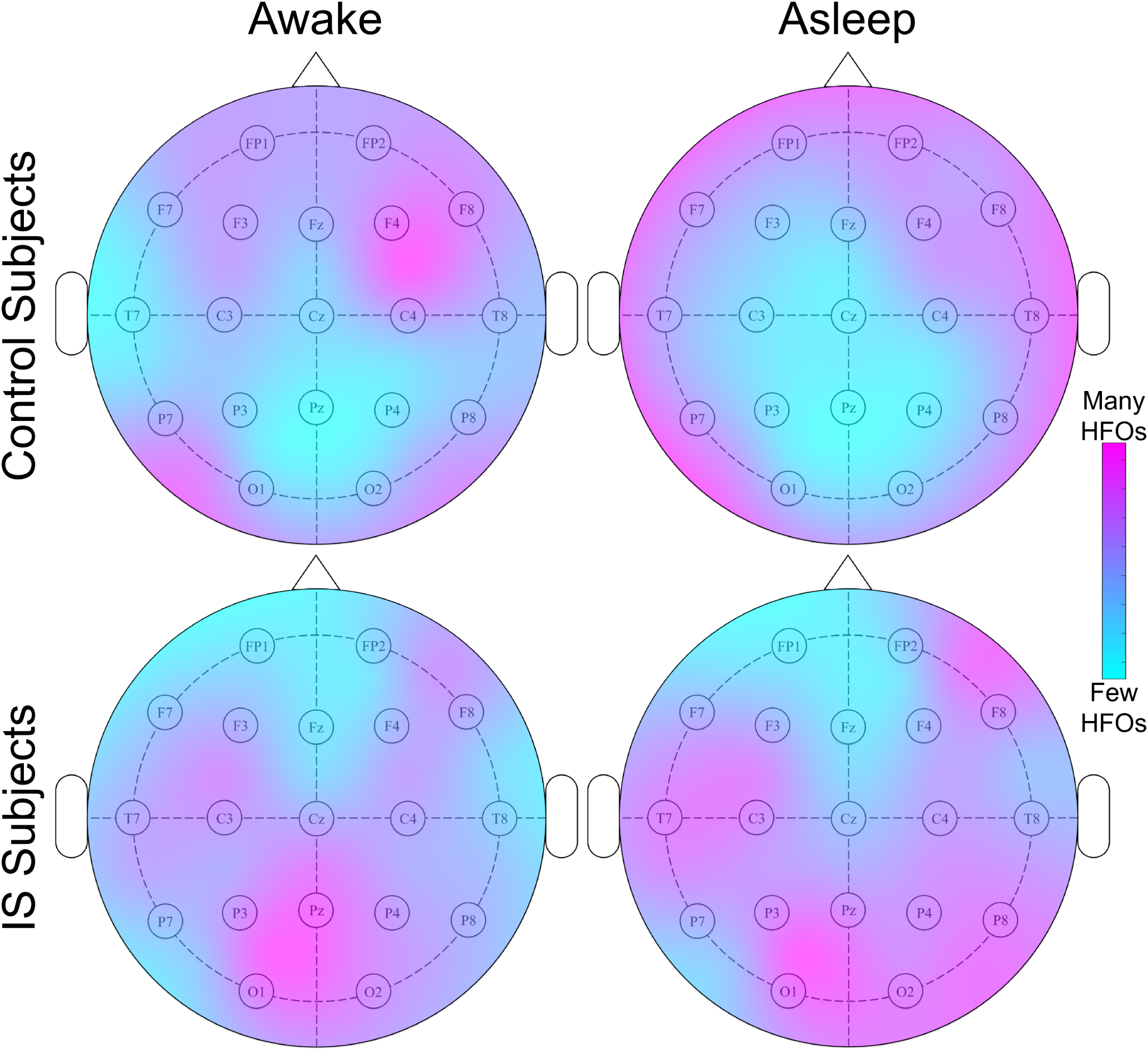
Median Spatial Distribution of HFOs Across Control and IS Subjects While Awake and Asleep. Areas in pink correspond to high HFO rates, and areas in cyan correspond to low HFO rates.

#### HFO Rates Before and After ACTH Initiation

Figure 5 depicts the HFO rates before ACTH initiation and >24 hours after the first dose of ACTH. In the 8 IS subjects included in this analysis, there was a significant decrease in HFO rates during sleep after ACTH was started (*p* =0.0078) per a Wilcoxon signed-rank test, with the median HFO rate across subjects decreasing from 6.8/min to 3.9/min. This was not true when patients were awake (*p* =0.11), with median HFO rate across subjects increasing from 2.5/min to 4.2/min. Recall that for this analysis, Subject 9 was excluded as she was initially diagnosed with focal epilepsy and started on phenobarbital, and Subjects 10, 12, and 13 were excluded because their pre- or post-ACTH study datasets were too small. Also note that only two out of the eight IS subjects included in this analysis responded clinically to ACTH treatment, as detailed below.

**Fig 5.**
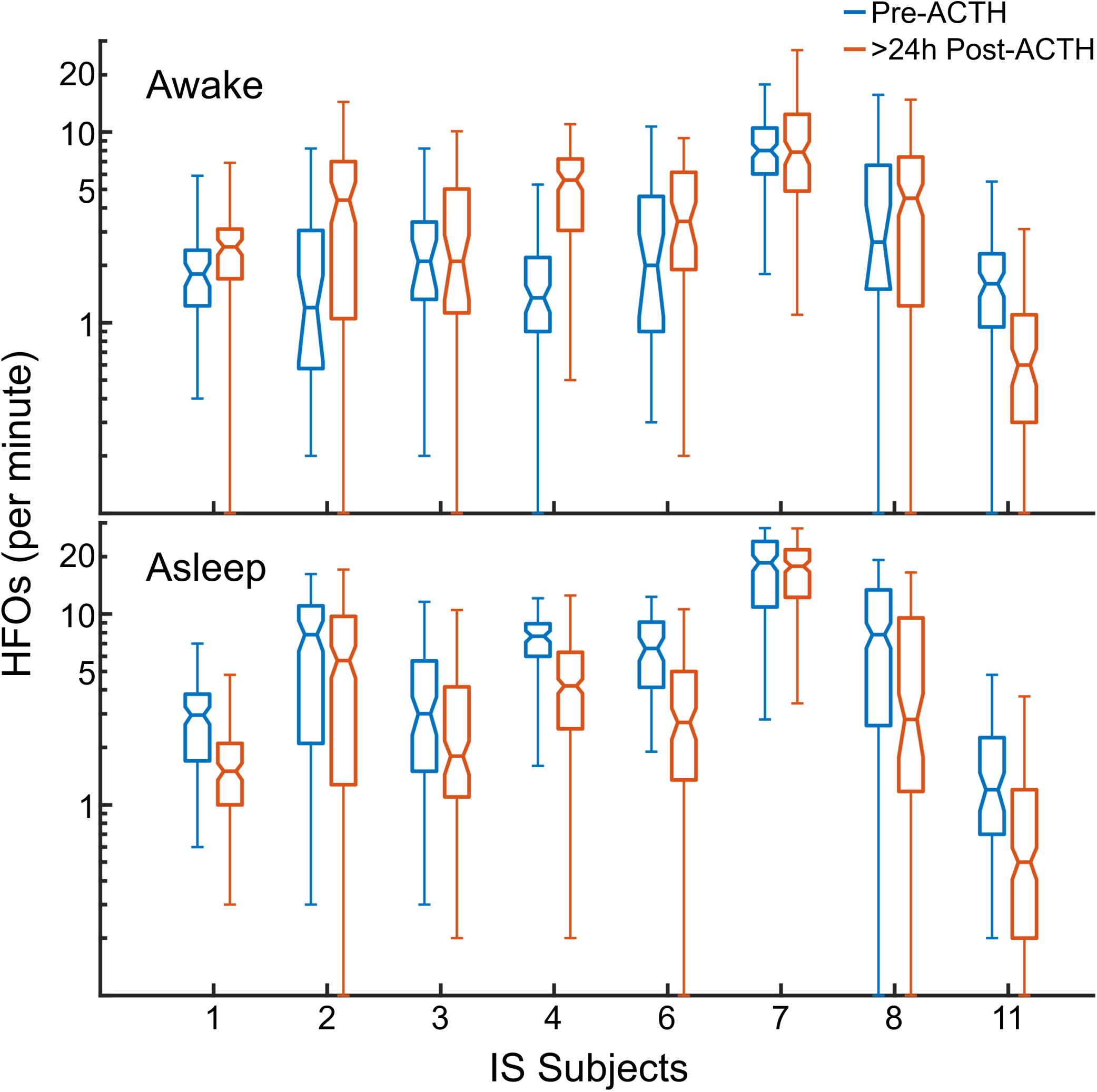
HFO Rates Before and After ACTH Initiation. HFOs during wakefulness (top) and sleep (bottom) before the start of ACTH therapy and at least 24 hours after it had been started were counted within multiple 10-minute intervals, and the distribution of HFO rates from these 10-minute intervals is depicted here. Note that a logarithmic y-axis scale is used for the box-and-whisker plots (interquartile range within box and all values within whiskers). Only IS Subjects underwent ACTH treatment during the study, and only eight out of thirteen had sufficiently large pre- and post-ACTH study datasets.

Five out of the thirteen IS subjects (namely Subjects 4, 8, 9, 12, and 13) initially responded clinically to ACTH therapy. The visual appearance of Subject 4’s EEG did not change substantially, although possibly contained slightly fewer discharges. She has had no evidence of IS relapse to-date, although she has had other non-IS-related seizures and her EEG continues to have discharges. Subject 8 exhibited moderate improvement in discharges on EEG by the end of the study. However, she relapsed about 6 months later and continues to have discharges on EEG. Subject 9 did exhibit improvement in discharges on her EEG with initial phenobarbital therapy for what was thought to be focal epilepsy. However, when she was diagnosed with IS approximately one month later, ACTH therapy was initiated and she responded well with no evidence of IS relapse to-date. Due to the short clinical and non-clinic recording duration for Subject 12, it was not possible to evaluate fully for EEG changes, although her subsequent EEG appears to have normalized, and we are unaware of any relapse. Subject 13’s recording duration was also too short to evaluate for visual changes, but she is known to have relapsed after subsequent ACTH wean. Note that all non-responders exhibited no substantial EEG changes after ACTH initiation except for Subject 6, who demonstrated improved spike frequency and amplitude by the end of the study, and Subject 10, who demonstrated more generalized EEG improvements by the end of the study.

## Discussion

This prospective study demonstrates fully automated detection of HFOs in long-term scalp EEG recordings and provides new insight into the characteristics of interictal scalp ripples in infants with IS. Patients with IS exhibited higher HFO rates than normal infant control subjects, and the rates were higher during sleep compared to wakefulness. In contrast, HFO rates for control subjects did not vary significantly across the sleep-wake cycle. Spatially, the distinction between the two groups was greatest in the central/posterior parasaggital region. HFOs in this region were frequently detected for IS subjects but rarely for control subjects, especially during sleep. ACTH therapy significantly and universally decreased HFOs in IS subjects during sleep, and the extent of this decrease tended to be large in subjects who clinically responded to treatment.

Our results are consistent with prior HFO studies using intracranial EEG. Based on these studies, it has become commonly accepted that interictal HFOs occur more frequently during sleep [55–57], although the difference is not always statistically significant [58, 59]. Bagshaw et al. [60] found similar diurnal HFO fluctuations in patients with focal epilepsy using implanted electrodes. The invasive nature of these measurements typically precludes analysis of data from controls that do not have epilepsy, but there are a few exceptions [61, 62].

Previous studies of interictal HFOs in scalp EEG relied on visual marking of events and exclusively used data collected during slow-wave sleep [26–28, 30, 31, 33, 38] or stage 2 sleep [32, 34]. This is the first report of HFO rates during wakefulness. In the most closely related study, Kobayashi et al. [38] found high rates of interictal scalp fast oscillations (41.0-140.6 Hz) in patients with West syndrome prior to treatment with ACTH. They reported a median ripple rate of approximately 7/minute in epilepsy patients and 0/min in healthy control subjects, which is similar to our median rates of 5.5/minute for IS patients and 0.98/minute for controls during sleep. They also found that ripples occurred most frequently in the P3-O1 and P4-O2 electrode pairs, consistent with our results. In that study, candidate HFOs were detected using automated time-frequency analysis, and the candidate events were visually reviewed to reject artifacts; 60 seconds of data were analyzed for each subject [38]. Given the significant differences in the amount of EEG data analyzed and the HFO detection methods, the consistency between our results and this prior study is noteworthy.

In any study of fast oscillations using scalp EEG, the influence of muscle artifacts is a concern. There is likely some basal level of muscle/ocular artifacts that escaped automated rejection in both the control and spasms subjects. However, these artifacts likely accounted for a small proportion of the total detected HFOs in the IS group, as this group exhibited substantially higher HFO rates during sleep when muscle/ocular movements (and their associated artifacts) are typically reduced. On the other hand, these artifacts may comprise a substantial proportion of the total detected HFOs in the control subjects, as HFOs were relatively more frequent during wakefulness in this group and were most prominent in artifact-prone EEG regions, e.g. the peripheral electrodes.

The occurrence of physiological ripples is an additional confounding factor. Physiological HFOs have been detected in the scalp EEG of healthy children, occurring primarily in the central and midline bipolar pairs [63, 64]. While rates varied widely across subjects, ripples occurred more frequently during light sleep and stage two sleep (stages N1 and N2), and approximately three-fourths of ripples co-occurred with sleep-specific transients [64]. Therefore, while false positive detections due to artifacts are less likely during sleep (due to reduced likelihood of muscle/ocular movements), there may be a simultaneous sleep-associated increase in true HFOs, leading to the overall lack of diurnal variation in the control subjects. This would likely occur in IS subjects, as well, but this effect appears to be masked by the overall higher rates of HFOs during sleep in this group.

Spatially, HFOs were very infrequent in the posterior parasagittal region (bipolar channels C3-P3, P3-O1, Fz-Cz, Cz-Pz, P4-O2, C4-P4) in the control subjects. In IS subjects, however, this was not the case, as a large portion of HFOs were detected from this region. The latter finding is consistent with a few prior studies [38, 65] that have suggested that interictal spikes and HFOs are predominantly phenomena of the posterior brain regions. It should be noted that in this study, the cortical areas of IS subjects with prominent discharges or hypsarrhythmia during EEG monitoring did not necessarily exhibit higher HFO rates. Further studies are needed to understand the electroneurophysiological etiology of these centro-posterior HFOs in IS patients, as well as the subgroup characteristics of the patients that exhibit this cortical phenomenon. Eventually, it may be possible to determine which patients are more or less likely to have infantile spasms based on the spatial distribution of their HFOs without the need for prolonged EEG monitoring.

Lastly, in the 8 IS subjects with sufficient pre- and post-ACTH data, we observed a significant decrease in HFO rates during sleep after the start of ACTH therapy. Note that while this decrease in HFO rate became apparent within days of starting ACTH therapy, it may also persist with continued treatment [38]. No similar decrease was appreciated for IS subjects during wakefulness, although it is possible that not enough time had passed to observe this effect. There was also an abnormally low number of responders to ACTH treatment in this prospective cohort (5 out of 13, with two of these relapsing after initial treatment), based on an assessment of the presence of spasms and hypsarrhythmia on the EEG after two weeks. The two responders (Subjects 4 and 8) included in the pre- and post-ACTH HFO rate analysis (Figure 5) showed large relative reductions in the HFO rates after starting ACTH therapy; however, this finding was not specific for responders and was also seen in non-responder Subjects 1, 6, and 11. Had we continued recording EEG for a longer period of time in patients that responded to treatment, we may even have expected to see a larger reduction in HFO rate. For example, Zijlmans et al. [23] reported a reduction in HFOs with medical therapy and an increase in HFO rates when medication was reduced. Current thinking is that effective treatment should eliminate EEG evidence of epileptic spasms [66] and epileptiform discharges [67]; however, it is unknown if it is necessary for pathological HFOs to be completely eliminated. Future studies are required to determine the long-term effects of ACTH therapy on HFOs in epileptic spasms patients. Many of the patients enrolled in this study have participated in serial longitudinal EEG monitoring at follow-up visits to help answer this question.

The current study has several limitations. Our analysis included only 13 subjects with IS, and a larger cohort would increase the power of the study. However, the subjects were prospectively recruited, and the large amount of data we analyzed for each subject increased the repeatability and robustness of the results. We did not perform visual validation of the automated HFO detections, which can be seen as either a strength (it is an unbiased approach and enables analysis of large amounts of data) or a weakness (our results undoubtedly contain some false positive detections due to muscle and other artifacts). The fact that the IS cohort contained many patients that did not respond to treatment hindered our ability to analyze the effect of ACTH on HFO rate. Also, we were not able to perform manual sleep staging for the IS subjects (unlike for the control subjects), due to the disruption in typical sleep patterns [48–51]. However, we expect our automated wake/sleep classifier to have performed sufficiently for the IS subjects’ datasets, given its 86% mean cross-validation accuracy for the control subjects and the assumption that sleep-wake features of EOG, EMG, ECG are consistent between control and IS subjects. Note that if the classifier performed at the level of random chance, we should expect no difference in the wakefulness/sleep characteristics, so as long as our classifier performed above chance level, the qualitative differences between wakefulness/sleep that were found in this study are real. In the future, it would also be useful to distinguish between the different stages of sleep for all subjects, as non-rapid eye movement (NREM) sleep is associated with higher rates of HFOs compared to REM sleep or wakefulness [57, 68, 69] and may be less prone to muscle and eye-movement artifacts [70]. An additional limitation was that we analyzed the entire long-term EEG recording without accounting for the possible presence of epileptic spasms. However, as these events are proportionally rare in longitudinal data, they are unlikely to have influenced the results. Lastly, we analyzed only HFOs in the ripple band (80-250 Hz), while there is evidence that fast ripples can also be detected in scalp EEG [31, 32]. Future studies should take steps to address these limitations.

Here we have presented an automated algorithm for HFO detection in scalp EEG that facilitates rapid analysis of large amounts of data in an unsupervised manner. This method is robust to the frequent occurrence of artifacts in the EEG, as we found significant differences between control and IS subjects in HFO rate, diurnal variation, and spatial distribution, as well as changes in HFO rates with antiepileptic therapy. These results support computational analysis of noninvasive scalp HFOs as a potentially valuable tool for the diagnosis and assessment of treatment response for epilepsy.

## References

1. Pavone P, Striano P, Falsaperla R, Pavone L, Ruggieri M. Infantile spasms syndrome, West syndrome and related phenotypes: What we know in 2013. Brain and Development. 2014; 36(9):739–751.

2. Hrachovy RA, Frost JD. Infantile epileptic encephalopathy with hypsarrhythmia (infantile spasms/West syndrome). Journal of clinical neurophysiology: official publication of the American Electroencephalographic Society. 2003;20(6):408–25.

3. Pellock JM, Hrachovy R, Shinnar S, Baram TZ, Bettis D, Dlugos DJ, et al. Infantile spasms: A U.S. consensus report. Epilepsia. 2010; 51(10):2175–2189.

4. Riikonen RS. Favourable prognostic factors with infantile spasms. European Journal of Paediatric Neurology. 2010; 14(1):13–18.

5. Hussain SA. Treatment of infantile spasms. Epilepsia Open. 2018; 3(S2):143–154.

6. Wirrell EC, Nickels KC. Electroclinical Syndromes: Infantile Onset. In: Swaimann KF, Ashwal S, Ferriero DM, Schor NF, Finkel RS, Gropman AL, et al., editors. Swaiman’s Pediatric Neurology. 6th ed. Elsevier; 2017. p. e1326–1345.

7. Lux AL, Osborne JP. A Proposal for Case Definitions and Outcome Measures in Studies of Infantile Spasms and West Syndrome: Consensus Statement of the West Delphi Group. Epilepsia. 2004; 45(11):1416–1428.

8. Caraballo RH, Fortini S, Reyes G, Carpio Ruiz A, Sanchez Fuentes SV, Ramos B. Epileptic spasms in clusters and associated syndromes other than West syndrome: A study of 48 patients. Epilepsy Research. 2016; 123:29–35.

9. Hrachovy RA, Frost JD, Kellaway P. Hypsarrhythmia: Variations on the Theme. Epilepsia. 1984; 25(3):317–325.

10. Donat JF, Lo WD. Asymmetric Hypsarrhythmia and Infantile Spasms in West Syndrome. Journal of Child Neurology. 1994; 9(3):290–296.

11. Hussain SA, Kwong G, Millichap JJ, Mytinger JR, Ryan N, Matsumoto JH, et al. Hypsarrhythmia assessment exhibits poor interrater reliability: A threat to clinical trial validity. Epilepsia. 2015; 56(1):77–81.

12. Mytinger JR, Hussain SA, Islam MP, Millichap JJ, Patel AD, Ryan NR, et al. Improving the inter-rater agreement of hypsarrhythmia using a simplified EEG grading scale for children with infantile spasms. Epilepsy Research. 2015; 116:93–98.

13. Crone NE, Sinai A, Korzeniewska A. High-frequency gamma oscillations and human brain mapping with electrocorticography. In: Progress in Brain Research. vol. 159; 2006. p. 275–295.

14. Smith MM, Weaver KE, Grabowski TJ, Rao RPN, Darvas F. Non-invasive detection of high gamma band activity during motor imagery. Frontiers in Human Neuroscience. 2014; 8.

15. Thürer B, Stockinger C, Focke A, Putze F, Schultz T, Stein T. Increased gamma band power during movement planning coincides with motor memory retrieval. NeuroImage. 2016; 125:172–181.

16. Zhang H, Fell J, Axmacher N. Electrophysiological mechanisms of human memory consolidation. Nature Communications. 2018; 9(1):4103.

17. Nowak M, Zich C, Stagg CJ. Motor Cortical Gamma Oscillations: What Have We Learnt and Where Are We Headed? Current Behavioral Neuroscience Reports. 2018; 5(2):136–142.

18. Kobayashi K, Oka M, Akiyama T, Inoue T, Abiru K, Ogino T, et al. Very Fast Rhythmic Activity on Scalp EEG Associated with Epileptic Spasms. Epilepsia. 2004; 45(5):488–496.

19. Iwatani Y, Kagitani-Shimono K, Tominaga K, Okinaga T, Kishima H, Kato A, et al. Ictal high-frequency oscillations on scalp EEG recordings in symptomatic West syndrome. Epilepsy Research. 2012; 102(1-2):60–70.

20. Kobayashi K, Miya K, Akiyama T, Endoh F, Oka M, Yoshinaga H, et al. Cortical contribution to scalp EEG gamma rhythms associated with epileptic spasms. Brain and Development. 2013; 35(8):762–770.

21. Jiruska P, Alvarado-Rojas C, Schevon CA, Staba R, Stacey W, Wendling F, et al. Update on the mechanisms and roles of high-frequency oscillations in seizures and epileptic disorders. Epilepsia. 2017; 58(8):1330–1339.

22. Wu JY, Koh S, Sankar R, Mathern GW. Paroxysmal fast activity: An interictal scalp EEG marker of epileptogenesis in children. Epilepsy Research. 2008; 82(1):99–106.

23. Zijlmans M, Jacobs J, Zelmann R, Dubeau F, Gotman J. High-frequency oscillations mirror disease activity in patients with epilepsy. Neurology. 2009; 72(11):979–986.

24. Andrade-Valenca LP, Dubeau F, Mari F, Zelmann R, Gotman J. Interictal scalp fast oscillations as a marker of the seizure onset zone. Neurology. 2011; 77(6):524–531.

25. Jacobs J, Zijlmans M, Zelmann R, Chatillon CÉ, Hall J, Olivier A, et al. High-frequency electroencephalographic oscillations correlate with outcome of epilepsy surgery. Annals of Neurology. 2010; 67(2):209–220.

26. Kuhnke N, Schwind J, Dümpelmann M, Mader M, Schulze-Bonhage A, Jacobs J. High Frequency Oscillations in the Ripple Band (80–250 Hz) in Scalp EEG: Higher Density of Electrodes Allows for Better Localization of the Seizure Onset Zone. Brain Topography. 2018; 31(6):1059–1072.

27. Melani F, Zelmann R, Dubeau F, Gotman J. Occurrence of scalp-fast oscillations among patients with different spiking rate and their role as epileptogenicity marker. Epilepsy Research. 2013; 106(3):345–356.

28. von Ellenrieder N, Andrade-Valença LP, Dubeau F, Gotman J. Automatic detection of fast oscillations (40–200Hz) in scalp EEG recordings. Clinical Neurophysiology. 2012; 123(4):670–680.

29. Zelmann R, Lina JM, Schulze-Bonhage A, Gotman J, Jacobs J. Scalp EEG is not a Blur: It Can See High Frequency Oscillations Although Their Generators are Small. Brain Topography. 2014; 27(5):683–704.

30. Kuhnke N, Klus C, Dümpelmann M, Schulze-Bonhage A, Jacobs J. Simultaneously recorded intracranial and scalp high frequency oscillations help identify patients with poor postsurgical seizure outcome. Clinical Neurophysiology. 2019; 130(1):128–137.

31. Pizzo F, Frauscher B, Ferrari-Marinho T, Amiri M, Dubeau F, Gotman J. Detectability of Fast Ripples (>250 Hz) on the Scalp EEG: A Proof-of-Principle Study with Subdermal Electrodes. Brain Topography. 2016; 29(3):358–367.

32. Bernardo D, Nariai H, Hussain SA, Sankar R, Salamon N, Krueger DA, et al. Visual and semi-automatic non-invasive detection of interictal fast ripples: A potential biomarker of epilepsy in children with tuberous sclerosis complex. Clinical Neurophysiology. 2018; 129(7):1458–1466.

33. Kramer MA, Ostrowski LM, Song DY, Thorn EL, Stoyell SM, Parnes M, et al. Scalp recorded spike ripples predict seizure risk in childhood epilepsy better than spikes. Brain. 2019; 142(5):1296–1309.

34. Gong P, Xue J, Qian P, Yang H, Liu X, Cai L, et al. Scalp-recorded high-frequency oscillations in childhood epileptic encephalopathy with continuous spike-and-wave during sleep with different etiologies. Brain and Development. 2018; 40(4):299–310.

35. Cao D, Chen Y, Liao J, Nariai H, Li L, Zhu Y, et al. Scalp EEG high frequency oscillations as a biomarker of treatment response in epileptic encephalopathy with continuous spike-and-wave during sleep (CSWS). Seizure. 2019; 71:151–157.

36. Kobayashi K, Yoshinaga H, Toda Y, Inoue T, Oka M, Ohtsuka Y. High-frequency oscillations in idiopathic partial epilepsy of childhood. Epilepsia. 2011; 52(10):1812–1819.

37. Kobayashi K, Ohuchi Y, Shibata T, Hanaoka Y, Akiyama M, Oka M, et al. Detection of fast (40–150 Hz) oscillations from the ictal scalp EEG data of myoclonic seizures in pediatric patients. Brain and Development. 2018; 40(5):397–405.

38. Kobayashi K, Akiyama T, Oka M, Endoh F, Yoshinaga H. A storm of fast (40-150Hz) oscillations during hypsarrhythmia in West syndrome. Annals of Neurology. 2015; 77(1):58–67.

39. Charupanit K, Nunez MD, Bernardo D, Bebin M, Krueger DA, Northrup H, et al. Automated Detection of High Frequency Oscillations in Human Scalp Electroencephalogram. In: 2018 40th Annual International Conference of the IEEE Engineering in Medicine and Biology Society (EMBC). IEEE; 2018. p. 3116–3119.

40. Nariai H, Beal J, Galanopoulou AS, Mowrey WB, Bickel S, Sogawa Y, et al. Scalp EEG Ictal gamma and beta activity during infantile spasms: Evidence of focality. Epilepsia. 2017; 58(5):882–892.

41. Kobayashi K, Akiyama T, Oka M, Endoh F, Yoshinaga H. Fast (40–150Hz) oscillations are associated with positive slow waves in the ictal EEGs of epileptic spasms in West syndrome. Brain and Development. 2016; 38(10):909–914.

42. Sparrow SS, Cicchetti DV, Saulnier CA. Vineland Adaptive Behavior Scales, Third Edition. 3rd ed. San Antonio, TX: Pearson; 2016.

43. Jasper HH. Report of the committee on methods of clinical examination in electroencephalography. Electroencephalography and Clinical Neurophysiology. 1958; 10(2):370–375.

44. Millett D, Stern JM. Fundamentals of EEG. In: Sirven JI, Stern JM, editors. Atlas of Video-EEG Monitoring. McGraw-Hill; 2011. p. 680.

45. Charupanit K, Lopour BA. A Simple Statistical Method for the Automatic Detection of Ripples in Human Intracranial EEG. Brain Topography. 2017; 30(6):724–738.

46. Burnos S, Hilfiker P, Sürücü O, Scholkmann F, Krayenbühl N, Grunwald T, et al. Human Intracranial High Frequency Oscillations (HFOs) Detected by Automatic Time-Frequency Analysis. PLoS ONE. 2014; 9(4):e94381.

47. Britton JW, Frey LC, Hopp JL, Korb P, Koubeissi MZ, Lievens WE, et al. Appendix 4. Common Artifacts During EEG Recording. In: Louis EKS, Frey LC, editors. Electroencephalography (EEG): An Introductory Text and Atlas of Normal and Abnormal Findings in Adults, Children, and Infants. Chicago: American Epilepsy Society; 2016.

48. Hrachovy RA, Frost JD, Kellaway P. Sleep characteristics in infantile spasms. Neurology. 1981; 31(6):688–688.

49. Kellaway P. Sleep and Epilepsy. Epilepsia. 1985; 26(s1):S15–S30.

50. Altunel A, Sever A, Altunel EÖ. Hypsarrhythmia paroxysm index: A tool for early prediction of infantile spasms. Epilepsy Research. 2015; 111:54–60.

51. Altunel A, Altunel EÖ, Sever A. The Utility of the Hypsarrhythmia Paroxysm Index and Sleep Spindles in EEG for Predicting Cognitive Outcomes in a Case Series of Infantile Spasms. Journal of Neurology & Neurophysiology. 2015; 06(05).

52. Fisher RA. The Use of Multiple Measurements in Taxonomic Problems. Annals of Eugenics. 1936; 7(2):179–188.

53. Mann HB, Whitney DR. On a Test of Whether one of Two Random Variables is Stochastically Larger than the Other. The Annals of Mathematical Statistics. 1947; 18(1):50–60.

54. Wilcoxon F. Individual Comparisons by Ranking Methods. Biometrics Bulletin. 1945; 1(6):80.

55. Dümpelmann M, Jacobs J, Schulze-Bonhage A. Temporal and spatial characteristics of high frequency oscillations as a new biomarker in epilepsy. Epilepsia. 2015; 56(2):197–206.

56. Clemens Z, Molle M, Eross L, Barsi P, Halasz P, Born J. Temporal coupling of parahippocampal ripples, sleep spindles and slow oscillations in humans. Brain. 2007; 130(11):2868–2878.

57. Staba RJ, Wilson CL, Bragin A, Jhung D, Fried I, Engel J. High-frequency oscillations recorded in human medial temporal lobe during sleep. Annals of Neurology. 2004; 56(1):108–115.

58. Schevon CA, Trevelyan AJ, Schroeder CE, Goodman RR, McKhann G, Emerson RG. Spatial characterization of interictal high frequency oscillations in epileptic neocortex. Brain. 2009; 132(11):3047–3059.

59. Liu S, Sha Z, Sencer A, Aydoseli A, Bebek N, Abosch A, et al. Exploring the time–frequency content of high frequency oscillations for automated identification of seizure onset zone in epilepsy. Journal of Neural Engineering. 2016; 13(2):026026.

60. Bagshaw AP, Jacobs J, LeVan P, Dubeau F, Gotman J. Effect of sleep stage on interictal high-frequency oscillations recorded from depth macroelectrodes in patients with focal epilepsy. Epilepsia. 2009; 50(4):617–628.

61. Gliske SV, Irwin ZT, Chestek C, Hegeman GL, Brinkmann B, Sagher O, et al. Variability in the location of high frequency oscillations during prolonged intracranial EEG recordings. Nature Communications. 2018; 9(1):2155.

62. Liu S, Gurses C, Sha Z, Quach MM, Sencer A, Bebek N, et al. Stereotyped high-frequency oscillations discriminate seizure onset zones and critical functional cortex in focal epilepsy. Brain. 2018; 141(3):713–730.

63. Mooij AH, Raijmann RCMA, Jansen FE, Braun KPJ, Zijlmans M. Physiological Ripples (±100 Hz) in Spike-Free Scalp EEGs of Children With and Without Epilepsy. Brain Topography. 2017; 30(6):739–746.

64. Mooij AH, Frauscher B, Goemans SAM, Huiskamp GJM, Braun KPJ, Zijlmans M. Ripples in scalp EEGs of children: co-occurrence with sleep-specific transients and occurrence across sleep stages. Sleep. 2018; 41(11).

65. Oka M, Kobayashi K, Akiyama T, Ogino T, Oka E. A study of spike-density on EEG in West syndrome. Brain and Development. 2004; 26(2):105–112.

66. Yamada K, Toribe Y, Kimizu T, Kimura S, Ikeda T, Mogami Y, et al. Predictive value of EEG findings at control of epileptic spasms for seizure relapse in patients with West syndrome. Seizure. 2014; 23(9):703–707.

67. Hayashi Y, Yoshinaga H, Akiyama T, Endoh F, Ohtsuka Y, Kobayashi K. Predictive factors for relapse of epileptic spasms after adrenocorticotropic hormone therapy in West syndrome. Brain and Development. 2016; 38(1):32–39.

68. Jirsch JD. High-frequency oscillations during human focal seizures. Brain. 2006; 129(6):1593–1608.

69. Sakuraba R, Iwasaki M, Okumura E, Jin K, Kakisaka Y, Kato K, et al. High frequency oscillations are less frequent but more specific to epileptogenicity during rapid eye movement sleep. Clinical Neurophysiology. 2016; 127(1):179–186.

70. von Ellenrieder N, Beltrachini L, Perucca P, Gotman J. Size of cortical generators of epileptic interictal events and visibility on scalp EEG. NeuroImage. 2014; 94:47–54.

